# Cortical grey matter mediates increases in model-based control and learning from positive feedback from adolescence to adulthood

**DOI:** 10.1101/2022.07.22.501096

**Authors:** Vanessa Scholz, Maria Waltmann, Nadine Herzog, Andrea Reiter, Annette Horstmann, Lorenz Deserno

## Abstract

Adolescents undergo maturation in cognition and brain structure. Model-based (MB) control is known to increase from childhood to young adulthood, which is mediated by cognitive abilities. Here, we asked two questions unaddressed in previous developmental studies: Firstly, what are the brain structural correlates of age-related increases in MB control? Secondly, how are age-related increases in MB control from adolescence to adulthood influenced by motivational context? A developmental sample (n=103, age: 12-42) completed structural MRI and an established task to capture MB control. The task was modified with respect to outcome valence by including (1) reward and punishment blocks to manipulate the motivational context and (2) an additional choice test to assess learning from positive vs. negative feedback. After replicating that an age-dependent increase in MB control is mediated by cognitive abilities, we demonstrate first-time evidence that grey matter density (GMD) in the parietal cortex mediates the increase of MB control with age. While motivational context did not relate to age-related changes in MB control, learning from positive feedback improved with age. Meanwhile, negative feedback learning showed no age effects. We present a first report that an age-related increase in learning from positive feedback was mediated by reduced GMD in the parietal, medial and dorsolateral prefrontal cortex. Our findings indicate that efficient brain maturation, as putatively reflected in lower GMD, in distinct and partially overlapping brain regions is a key developmental step towards age-related increases in planning and value-based choice.

**Significance Statement:** Adolescents undergo extensive maturation in cognition and brain structure. Interestingly, model-based decision-making is also known to increase from childhood to adulthood. Here, we demonstrate for the first time that grey matter density in the parietal cortex mediates an age-dependent increase in model-based control. An age-related increase in positive feedback learning was mediated by reduced grey matter density in the parietal, medial and dorsolateral prefrontal cortex. Interestingly, a manipulation of motivational context (gain reward vs. avoid punishment) did not impact age-related changes in model-based control. These findings highlight that efficient brain maturation in distinct and overlapping cortical brain regions constitutes a key developmental step towards increases in model-based planning and value-based choice.

## Introduction

Value-based learning and decision-making are guided by model-based and model-free reinforcement learning (RL) systems (Daw et al., 2005, 2011; Daw & Dayan, 2014). The model-based (MB) system relies on a model of the environment by mapping states, actions and outcomes in a probabilistic manner (Daw et al., 2005, 2011; Daw & Dayan, 2014; Dayan & Niv, 2008). This enables flexible behavior but is cognitively demanding. MB contributions to control were shown to increase from childhood into young adulthood (Bolenz et al., 2017, 2019; Decker et al., 2016; Nussenbaum et al., 2020; Vaghi et al., 2020), which was mediated by cognitive abilities (Nussenbaum et al., 2020; Potter et al., 2017). Our present study replicates these findings and examines hitherto unaddressed research questions. Firstly, what are the neuroanatomical correlates of age-related increases in MB control? Secondly, are these increases distinctively influenced by motivational context? Finally, does learning from positive and negative feedback change with age?

After a marked increase of gray matter (GM) from infancy to childhood (Gilmore et al., 2012; Knickmeyer et al., 2008), the adolescent brain shows a profound GM reduction in frontal, parietal and temporal cortices (Blakemore, 2012; Tamnes et al., 2013; Ziegler et al., 2019), potentially from synaptic pruning (Giedd et al, 1999, Gogtay et al. 2004, Sowell et al. 2001). This may lead to a more efficient brain organization, including myelination (Fuhrmann et al., 2015), hypothetically underlying the coinciding improvement in cognition, such as working memory (WM) (Bunge & Wright, 2007; Jolles et al., 2011; Tamnes et al., 2013). In previous work, MB control was positively related to cognitive abilities including WM (Eppinger et al., 2013; Otto et al., 2013) and processing speed (Reiter et al., 2016; Schad et al., 2014) and increasing MB control with age was mediated by cognitive abilities (Nussenbaum et al., 2020; Potter et al., 2017). GM in dorsolateral and ventromedial prefrontal cortex correlated with MB control in adults (Deserno et al., 2015; Voon, Derbyshire, et al., 2015) and prefrontal and parietal cortices were shown to encode state predictions, a neural signature of MB RL (Gläscher et al., 2010). However, it remains unclear whether structural brain maturation indeed mediates increases in MB control from adolescence to adulthood.

Previous work (Cauffman et al., 2010; Cohen, 2011; Silverman et al., 2015; Van Den Bos et al., 2012; Van Leijenhorst et al., 2010) also suggested that effects of outcome valence on RL (Daw & Dayan, 2014; Sutton & Barto, 1998) may change across development, by showing elevated reactivity in adolescents towards rewards overall or relative to punishment (but see Nussenbaum & Hartley, 2019; Rosenbaum et al., 2022). Thus, adolescents may be more willing to exert cognitive resources, i.e. employ MB control to gain rewards relative to punishment avoidance. However, the only study examining such effects on MB control found no age-dependency of contextual valence on MB control (Bolenz et al. 2021). Meanwhile, a large body of work links differences in positive vs negative feedback learning to positive and negative reward prediction errors (RPEs) (Frank et al., 2004, 2005, 2007). Phasic dopamine responses to RPE are asymmetric such that bursts for positive RPEs are larger than dips for negative RPEs (Montague et al., 1996; Schultz et al., 1997). In adolescence, RPE signaling in the ventral striatum is enhanced compared to adults (Cohen et al. 2010). An established test capturing this asymmetry by comparing positive versus negative feedback learning is derived from a probabilistic selection task (Frank et al., 2004) but has not yet been applied in a developmental sample.

Thus, we aimed to experimentally separate and study two developmentally relevant aspects of outcome valence: (1) learning in a reward or punishment (motivational) context and (2) learning from positive vs. negative feedback. For this, we recruited a developmental sample that completed structural neuroimaging and a modified “2step” task (Doll et al., 2016; Voon, Baek, et al., 2015), to study the structural correlates of the development of MB control for reward and punishment and learning from positive vs. negative feedback.

## Material and Methods

### Sample

A developmental sample of n = 103 participants (age range 12-50 years, 48 = female, 55 = male) was recruited as part of a larger study. This subsample was specifically screened to exclude current mental health diagnosis (also see pre-registered study protocol at <u>10.17605/OSF.IO/FYN6Q</u>). Participation consisted of two appointments. On day one, each participant completed a modified sequential decision-making task, the 2-step task (Daw et al., 2011) behaviorally, which was followed by a choice test capturing learning from positive vs. negative feedback (Doll et al., 2009, 2011, 2016; Frank et al., 2004) after a 30 minute break, during which participants either completed another task or questionnaires. This was all part of a larger task battery. They also underwent a battery of cognitive measures [Digit Symbol Substitution Test (DSST) for processing speed (Wechsler, 1997), Digit Span for working memory, (Wechsler, 1997), trail making test (TMT) A and B (Army Individual Test Battery, 1944) for executive functioning and a German vocabulary test (Lehrl, 2005)] (see study protocol for details). On the second day, participants underwent structural MRI. Participants were reimbursed with 9 Euro per hour for participation and a bonus payment based on task performance. The study was in agreement with the declaration of Helsinki and approved by the ethics board of the medical faculty at the University of Leipzig (385/17-ek). All participants were informed about study proceedings and gave informed consent before participation.

### Experimental tasks

The study included a sequential decision-making task, which encompassed two major modifications to address the research questions outlined in the introduction. These changes included: (1) separate reward and punishment blocks during the task (Voon, Baek, et al., 2015) to test effects of motivational context on MB control; (2) a choice test akin to the probabilistic selection task (Frank et al., 2004), following the 2-step task (Doll et al., 2016) testing learning from positive vs. negative feedback.

#### Sequential decision-making task

Similar to Daw et al., 2011, participants were presented with two different cue pairs in the first stage and had to select one cue to continue to the second stage (Fig. 1). Each cue was associated with a probabilistic transition to one of the two second stage states, a common transition of 70%, and a rare transition of 30%. Transition probabilities were fixed across the task. At the second stage, participants again had to choose between two cues and received an outcome (reward, neutral outcome, punishment, depending on the within-subject manipulation of motivational context, see below). Outcome probabilities changed slowly but constantly over time according to a Gaussian random walk. Thus, to maximize outcome in this task, participants need to track these continuously changing outcome probabilities. Importantly, participants were explicitly told that the task’s transition structure (common/ rare) would be constant over the task and that one transition was going to be more probable than the other. They also underwent practice trials to ensure that they had understood the task. By making use of knowledge about the transition structure, individuals could exert of MB control over choices. We employed a task variant, which included an additional within-subject manipulation of motivational context for positive and negative outcomes (Voon, Baek, et al., 2015; Worbe et al., 2016). Each participants completed 2 task blocks (each 100 trials). The reward block had monetary rewards displayed on the screen (+ 20 cent) or neutral feedback, while the punishment block used punishments (−20 cent) alongside neutral feedback. Block order was randomized across participants to avoid order effects. Participants were instructed to maximize outcome and could do so by learning to select the currently most rewarded cue in the reward block or avoiding negative outcomes in the punishment block. They were told that they would be paid out a bonus dependent on the rewards gained during the experiment.

**Figure 1:**
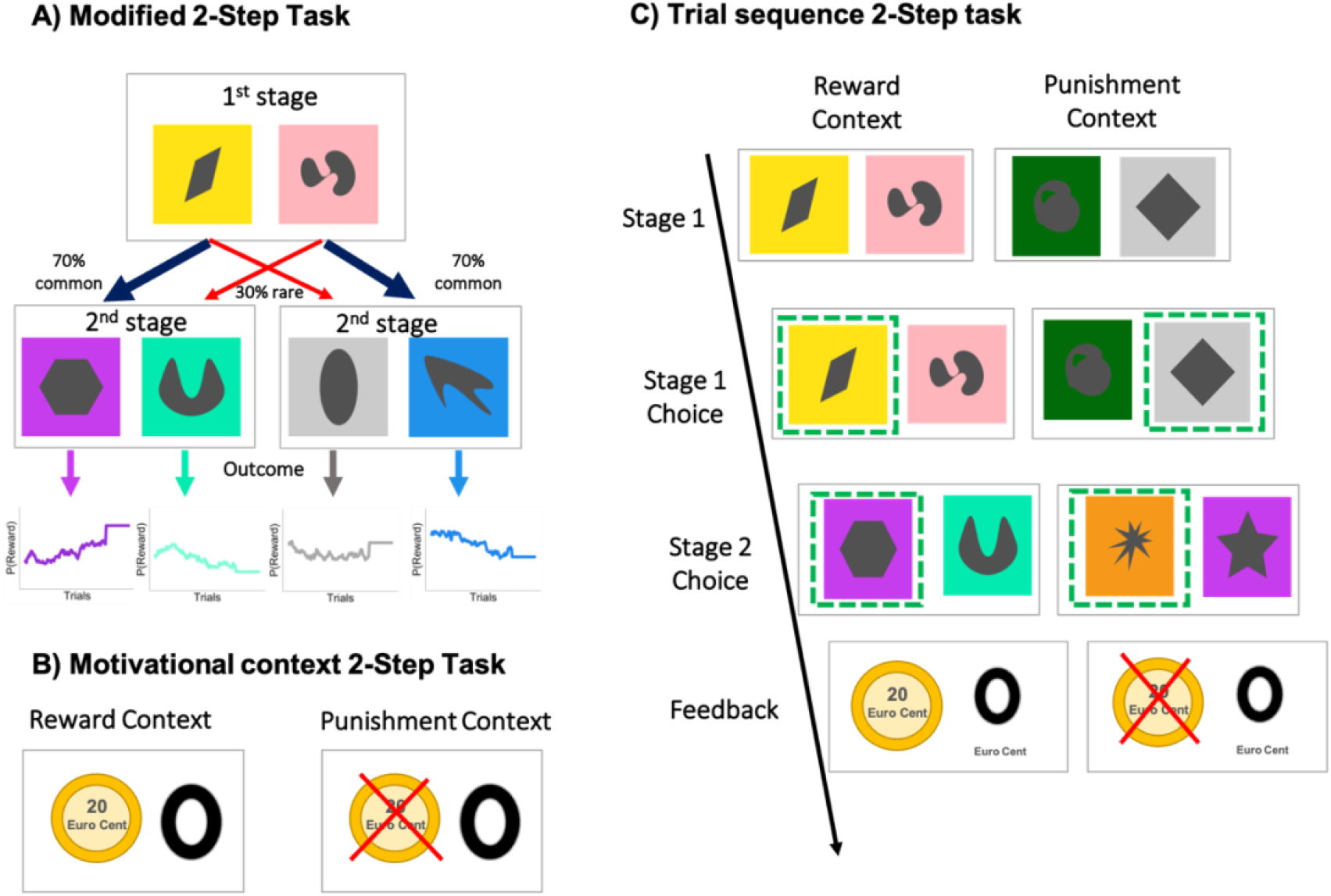
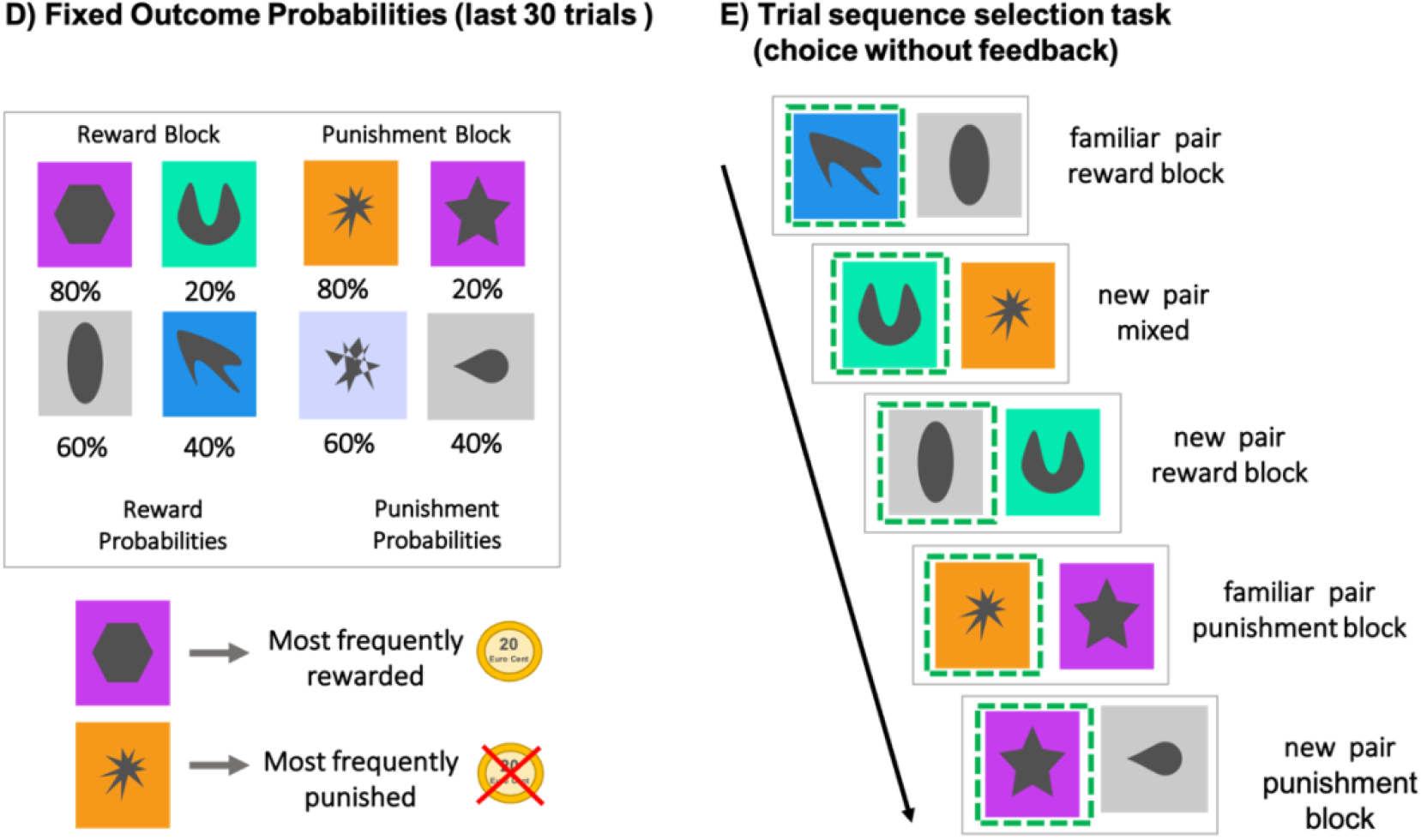
Task set-up sequential decision-making task. In the sequential decision-making task, a first stage choice led to one of two possible second stages, during which a second choice had to be made. After this second stage choice, participants received a reward or neutral outcome (rewards were replaced by punishments in the punishment context). The probability of receiving a reward or punishment was determined by continuously changing probabilities, i.e. Gaussian random walks. Transition probabilities from stage 1 to stage 2 were fixed and are either considered common (70%) or rare (30%) **B) Depiction of motivational context**. Outcome options for the reward and punishment context **C) Trial sequence sequential decision-making task**, presented once for the reward context which employs positive and neutral outcomes, and once for the punishment context. **D) For the choice test: fixing reward probabilities in the last 30 trials:** Here the fixed outcome probabilities assigned to the two familiar pairings for the last 30 trials of each motivational context in the sequential decision-making task are depicted. E**) Overview of choice phase** depicting cue selection across five trials without feedback to make sure that no further feedback-based learning occurs. Participants were required to select what they thought was the better cue from either familiar or recombined, new pairs.

#### Choice test

To examine age-dependent differences in learning from positive and negative outcomes and the impact of motivational context on learning, we employed a variant of a previously established probabilistic choice task (Doll et al., 2009, 2011, 2016; Frank et al., 2004) (see Figure 1E). To enable a choice test subsequent to the 2step task, reward probabilities remained stable for the last 30 trials in each block (reward/ punishment) of the 2step task. Hence, the previously slowly changing reward probabilities of one 2^nd^ stage pair were fixed to 80% : 20% chance of winning (or losing for the punishment block), whereas the probabilities were fixed at 60% : 40% for the other second stage pair. Thus, in the last 30 trials of each block, participants could learn the cue values in a stable manner (e.g. infer the most frequently rewarded and least frequently punished cues) (see Figure 1D). In the ensuing choice task, participants were presented with 28 different cue pairs, which were presented 3 times amounting to a total of 84 trials. These consisted of 4 familiar pairs they had previously encountered during the second stage of the 2-step task and 24 unfamiliar pairs from newly combined cues from the second stage. Unfamiliar pairs could be grouped into two categories 1) two recombined cues derived from the same motivational context (reward or punishment) (8 pairs) and 2) mixed pairs combining cues from the reward and punishment block (16 pairs). Of note, mixed pairs represent a unique feature of this task version and have not been employed frequently in previous work (Palminteri et al., 2016). They were introduced to increase the general level of task difficulty and variance of performance thus allowing better assessment of inter-individual differences across participants. Participants were instructed to select the ‘best’ cue from each pair upon presentation and unlike before, they did not receive feedback after having made their choice to disable further feedback-based learning. Thus, the selection task examines the values participants have learned for each cue throughout the previous task phase, i.e. the value they encoded for the respective stimulus. Of note, participants were aware of a performance-based bonus payment related to both tasks.

#### Analysis of 2-step task

To examine which factors impacted first-stage choice behaviour on the subsequent trial, we computed linear mixed effects models using the lme4 package implemented in R, (http://cran.us.r-project.org) with the optimizer bobyqa and the maximal number of iterations set to n = 1e+9. The model included participants’ trial-by-trial first-stage choices (stay or switch in a given trial n as compared to the previous trial n-1) as dependent variable (DV). Second stage feedback (positive vs. negative; in the reward block this refers to positive vs neutral outcome while in the punishment block this refers to neutral vs negative outcome) and transition type (rare vs. common) from the previous trial and motivational context (reward vs. punishment block) as within-subject fixed effects in the model. The model was estimated with a full random effects structure, i.e. all fixed effects were included as random effects:

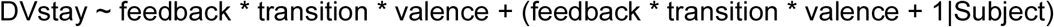

To determine whether we could replicate age-dependent changes in MB control as reported by Decker and colleagues (2016), we extracted the individual slope of the fixed effects interaction term feedback x transition as a valence-dependent individual estimate of MB control and correlated it with age. To determine age-dependent changes in MB control as a function of motivational context, we ran the model separately for the reward and punishment blocks and correlated the extracted estimates of MB control _reward_ and MB control _punishment_ with age, respectively. Given non-normal distributions of some variables, we assessed correlations using Spearman correlation coefficients. We excluded one participant due to missing data (n = 30 fewer trials on task) and a second participant who was an age outlier with respect to the remaining sample (8-year age gap, age 50). Thus, from the initial sample of n =103, a final sample of n = 101 remained for behavioral analysis after exclusion (M:F: 54:47, mean age = 23.03 (7.98), age range: 12-42).

#### Cognitive measures and MB control

We also examined the replicability of previous findings (Schad et al., 2014) showing a link between cognition and MB control. For this, we ran mediation analyses using the mediation package (Tingley et al., 2014) implemented in R to test whether the association between age and MB control was mediated by cognitive abilities. We used nonparametric bootstrapping with 10000 simulations to determine the average causal mediation effect between age and MB control mediated by cognitive measure.

#### Analysis of choice test

Due to fewer participants completing this task and one participant with a missing response rate of > 95% on the selection task only, a sample of n = 90 was available for behavioral data analysis of the selection task. We studied participants’ tendency to learn from positive and negative feedback using mixed-effects modelling and assessed age effects using correlational analysis. Here, we examined the difference between selecting the best cue for reward or punishment pairs (choose 80% rewarded or 20% punished over all cues) relative to avoiding the worst cue in reward or punishment pairs (avoid the cue that was 20% rewarded or 80% punished). This difference captures the shift towards learning more from positive or negative feedback, while no difference indicates equal learning from both modalities. We restricted analysis to reward pairs including the most frequent winner (80% positive outcome) or the least frequent winner (20%) and punishment pairs with the least frequent loser (20% negative outcome) or the most frequently punished cue (80%). Selecting the best cue, previously termed cue “A”, represents an individual’s tendency to learn from positive feedback, while avoiding the worst cue, also known as “B” from previous work, is said to capture learning from negative feedback (Doll et al., 2016; Frank et al., 2004; Waltz et al., 2007). The following equation was employed to assess the impact of learning from positive and negative feedback on optimal decision-making:

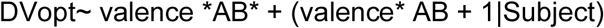

We concluded this analysis step by assessing whether MB control was linked to learning from positive or negative feedback in the selection task using correlational analysis.

### Structural brain data

#### Data acquisition structural imaging

Whole-brain T1 weighted image acquisition took place on a 3T Magnetom Skyra scanner (Siemens Healthcare, Erlangen, Germany). Structural data for subsequent VBM analysis was acquired with an echo time (TE) of 2,98 ms, a repetition time (TR) of 2300 ms, voxel size: 1.0× 1.0 × 1.0 mm, a field of view (FOV) of 256 mm, 176 slices with a slice thickness of 1 mm. For n =16 participants a different multi-echo sequence with the following parameters was employed: Echo time (TE) of [1,96; 5,83; 8,78; 11,73; 15,18] ms, a repetition time (TR) of 7000 ms, voxel size: 1.0× 1.0 × 1.0 mm, a field of view (FOV) of 256 mm, 192 slices with slice thickness of 1 mm.

#### Preprocessing of structural brain data

To examine morphometrical changes in the brain’s grey matter density (GMD), structural imaging data was preprocessed and analyzed using the statistical parametric mapping software (SPM) 12 (http://www.fil.ion.ucl.ac.uk/spm/software/spm12) and the Computational Anatomy Toolbox (CAT12) (http://dbm.neuro.uni-jena.de/cat) Version 12.7 (1728). Preprocessing followed the default pipeline outlined in the CAT12 manual and encompassed normalization to a template space, tissue segmentation into GM, white matter (WM) and cerebrospinal fluid (CSF), estimation of total intracranial volume (TIV) and smoothing. Smoothing was accomplished using an 8 mm full-width at half-maximum (FWHM) kernel (for more information see CAT12 manual). A 0.1 absolute masking threshold was applied to the data. Before analysis, data was screened and the weighted average image quality ratings implemented in CAT12 was deemed satisfactory. TIV was then estimated for the entire sample. Evaluating design-orthogonality using SPM provided evidence of a non-orthogonality between the TIV regressor and our main effects of interest, namely MB-control and learning from positive feedback. Hence, TIV was not included as regressor in our GLMs. Instead, we used global scaling to scale the data based on individual TIV values thereby avoiding the removal of variance of interest (see manual). Of note, all GLMs also included an additional regressor of no interest coding for the scanning sequence.

#### Statistical analysis of GM, MB control and age

A sample of n = 98 was available for MRI analysis, as 3 participants did not undergo scanning. We used two GLMs to assess effects of age and MB control on GMD. The first model (GLM_MBC_) comprised a regressor with the extracted individual slopes for MB control extracted from the mixed model described above to examine any changes in GMD that were associated with MB control in general. Meanwhile, our second model included regressors for age, MB control and their interaction (GLM_MBC_Full_) to determine which changes in GMD that were linked to MB control would remain when controlling for age. The interaction term of this model additionally allowed us to assess whether certain changes in GMD differed as a function of age and MB control.

#### Statistical analysis of GM and feedback learning

A sample of n = 88 was available for the structural brain analysis after MRI and selection task drop out. To probe the association between GMD and selection task performance, we assessed two more models. The first model comprised the extent to which participants had learned from positive feedback, i.e. had selected the better cue in a newly combined stimulus pair, as results had shown age-dependent changes for positive feedback learning only (GLM_PosFB_). This was again done to inspect which GMD changes were generally linked to positive feedback learning. The second model comprised 3 regressors: Age, “A”/ better cue selected and the interaction (GLM_PosFB_Full_). This model allowed us to isolate changes in GMD related to positive feedback learning independent of age effects, to assess age-dependent GMD changes and to determine whether GMD changes differed as a function of age and feedback learning. We chose to examine positive feedback learning only, as positive feedback had shown a strong age effect unlike negative feedback learning.

#### Regions of interest (ROI)

Results of all brain structural analyses were examined 1) using FWE correction of peak levels for multiple comparison on a whole-brain level and 2) using small-volume correction relying on three a-priori defined masks that were derived from the Automated Anatomic Labeling (Tzourio-Mazoyer et al., 2002). This included the vmPFC (superior medial frontal and medial orbital gyrus), dorsolateral PFC (dlPFC) (middle frontal gyrus) and the parietal cortex (inferior parietal gyrus and angular gyrus) motivated by previous accords of the involvement of those regions in MB control in fMRI studies using a similar task (Daw et al., 2011; Deserno et al., 2015; Gläscher et al., 2010; Voon, Derbyshire, et al., 2015). Results were considered significant with a p-value < 0.02 (0.05/3 for three regions of interest) to correct for multiple comparisons.

#### Mediation analysis

Given the lack of previous work examining the structural correlates of age-dependent changes of MB-control and feedback-learning, we relied on mediation analysis to examine whether maturational changes in GMD mediated the association between age and positive feedback learning. For this, we employed age as independent variable (IV), MB control (or positive feedback learning) as DV and GMD as mediator.

To assess whether certain GMD changes mediated the association between age and MB control or age and learning from positive feedback, we used the previously computed GLM with a regressor for MB control and a second GLM including age as regressor. For both GLMs, we created an F-contrast assessing any associated changes in either direction for the regressors age and MB control respectively. Each statistical map was thresholded at p =.001 and cluster size ≥ 20 voxels at the whole-brain level and subsequently exported as a mask image. We then created a conjunction mask from the two F contrast masks and the parietal cortex mask, for which results had survived SVC for the GLM_MBC_. We then extracted GMD estimates from this mask region using the get_totals script by Ged Ridgway (http://www.cs.ucl.ac.uk/staff/G.Ridgway/vbm/get_totals.m). Of note, as no clusters in the vmPFC or dlPFC were significantly associated with MB control, conjunction masks did not yield a common region from which to extract GMD estimates. For mediation analysis, we relied on the mediation package in R.

To follow up on age-dependent effect for feedback learning, we employed the same approach as outlined for the mediation analysis including MB control to analyze mediation effects related to feedback learning. The only difference being, that we now based the conjunction mask on the GLM for (positive) feedback learning. Of note, as we only found age-dependent effects of learning from positive feedback, this analysis focused on positive feedback learning. Again, we examined the mediating role of GMD derived from the vmPFC, dlPFC and parietal cortex conjunction masks.

## Results

### Replication: MB control increases with age

Using correlational analysis, as done by Decker et al (2016), we replicated age-dependent increases of MB control (r_s_(99)= 0.31, p-value = .002) (Decker et al., 2016; Nussenbaum et al., 2020) (Figure 2). Also, MB control in the punishment and reward block both significantly and comparably correlated with age (MB control _Punishment_ : r_s_(99)= 0.28, p = .004; MB control _Reward_ : r_s_(99)= 0.27, p = .007) indicating that age-dependent improvements in MB control did not differ as a function of motivational context (reward gain vs punishment avoidance blocks). This replicates the previously reported absence of general effects of motivational context (Bolenz & Eppinger, 2021)

**Figure 2.**
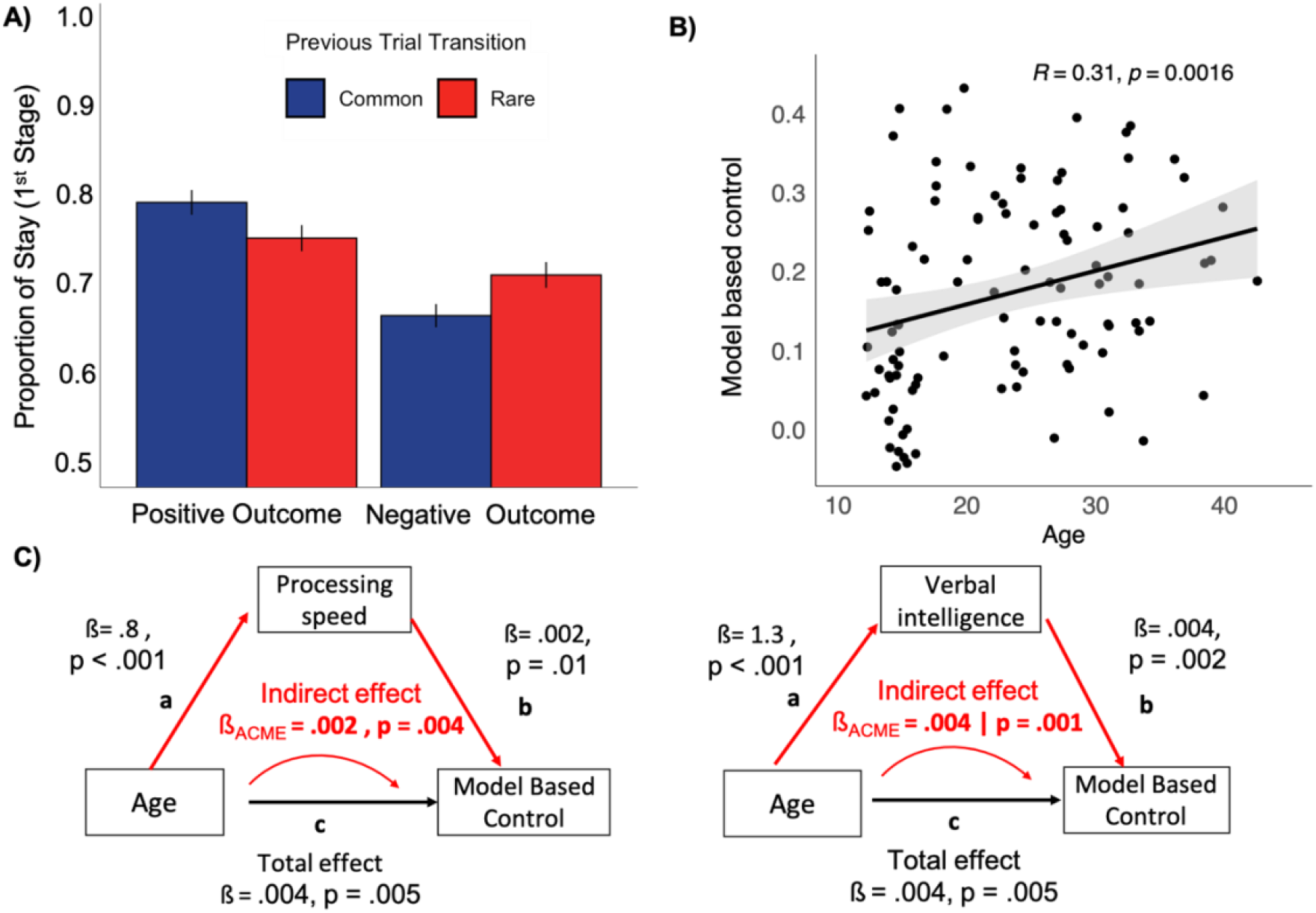
**A)** Depiction of task performance overall. **B)** Association between model-based control with age. **C)** Mediation effects of the relationship between age and model-based control through cognitive abilities, namely processing speed as measured by the DSST and verbal intelligence indexed through MWTB scores. P-values below .05 are considered significant.

### Replication: Cognition mediates age-related increases in MB control

Mediation analysis provided evidence of a significant partial mediation of processing speed measured using the *DSST*, accounting for 42,1 % (p = .006) of the total effect of age on MB control (Indirect effect = 0.002, CI [0.0005 – 0.003], p = .004; direct effect = 0.003, CI [-0.0007 – 0.01], p = .1; Total effect = 0.004, CI [0.002 – 0.01], p = .006). For verbal intelligence, mediation analysis suggested a full mediation for the effect of age on MB control (p = .005) (Indirect effect = 0.005, CI [0.002 – 0.01], p = 0.002; Direct effect = -0.0001, CI [-0.004 – 0.002], p-value = .98; total effect = 0.004, CI [0.002 – 0.01], p = .003) (see Figure 2). Both effects were significant after multiple comparison correction (α = 0.05/6 = 0.008). No significant mediation effects were observed for working memory, short term memory capacity, visual attention and general executive functioning (all p-values > 0.05).

### Structural brain correlates of MB control

For the GLM_MBC_, including individual estimates of MB control and a covariate controlling for the structural sequence, no significant association for MB control with GMD survived FWE correction on the whole brain level. Using small volume correction in our *a-priori* defined regions of interest, we found GMD in the parietal cortex to be associated with MB control (MNI Peak coordinate: -48 -50 57, k= 494, T = 4.42, p_FWE Peak corr_ = .008), which did not reach significance for the effect of mb control in dlPFC GMD (MNI Peak coordinate: -32 57 2, k= 10, T = 3.29, p_FWE Peak corr_ = .3). In the vmPFC no suprathreshold clusters were identified. In the GLM_MBC_Full_, age negatively correlated with GMD across the entire cortex, most prominently for a large cluster comprising the right frontal superior gyrus, the left temporal middle gyrus and the right supramarginal gyrus (MNI Peak coordinate: 18 56 16, k= 37120, T = 6.02, p_FWE Peak corr_ = .001). We did not find significant association with GMD for MB control, while controlling for the effects of age, nor for the interaction for age x MB-control in this whole-brain analysis (pFWE _**corr peak/ cluster**_ > .05). Likewise, we did not find any significant clusters using SVC for our three pre-defined regions for the MB control regressor or the interaction regressor.

Next, we examined whether GMD in parietal regions mediated the association between age and MB control. In a mediation model, the association between age and MB control was partially mediated by GMD in the parietal cortex, with GMD accounting for 67.7% of the total effect of the relationship between age and MB control (p = .007) (see Figure 4).

**Fig. 3.**
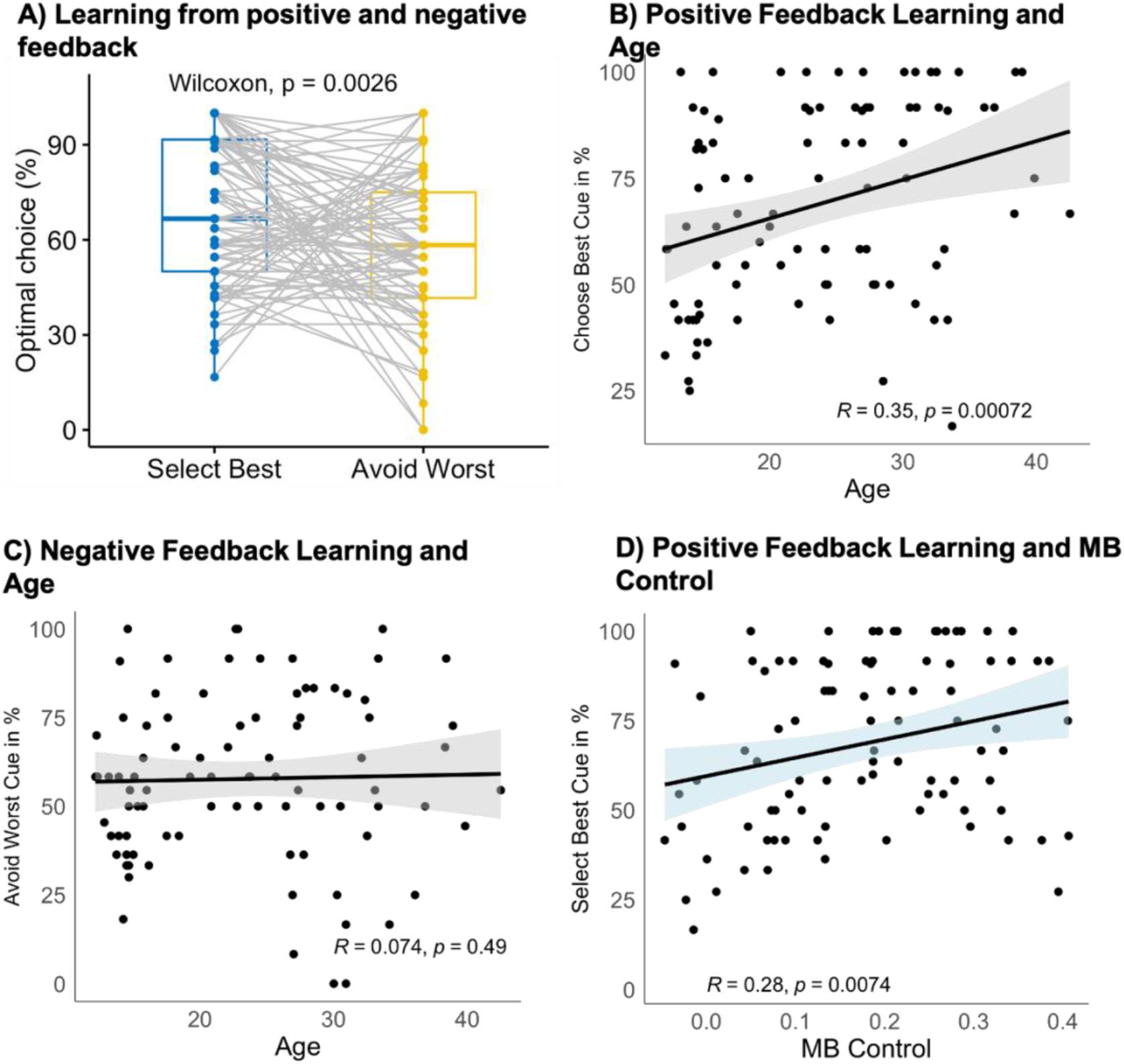
A) Overview of participants’ performance when choosing the best cue and avoiding the worst cue across both blocks. Participants show better performance when choosing the best cue, i.e. they learn better from positive feedback relative to learning from negative feedback or avoiding the worst cue across blocks. B) Age dependent increases of learning from positive feedback. C) Learning from negative feedback did not show age effects D) MB control was positively associated with positive feedback learning.

**Figure 4.**
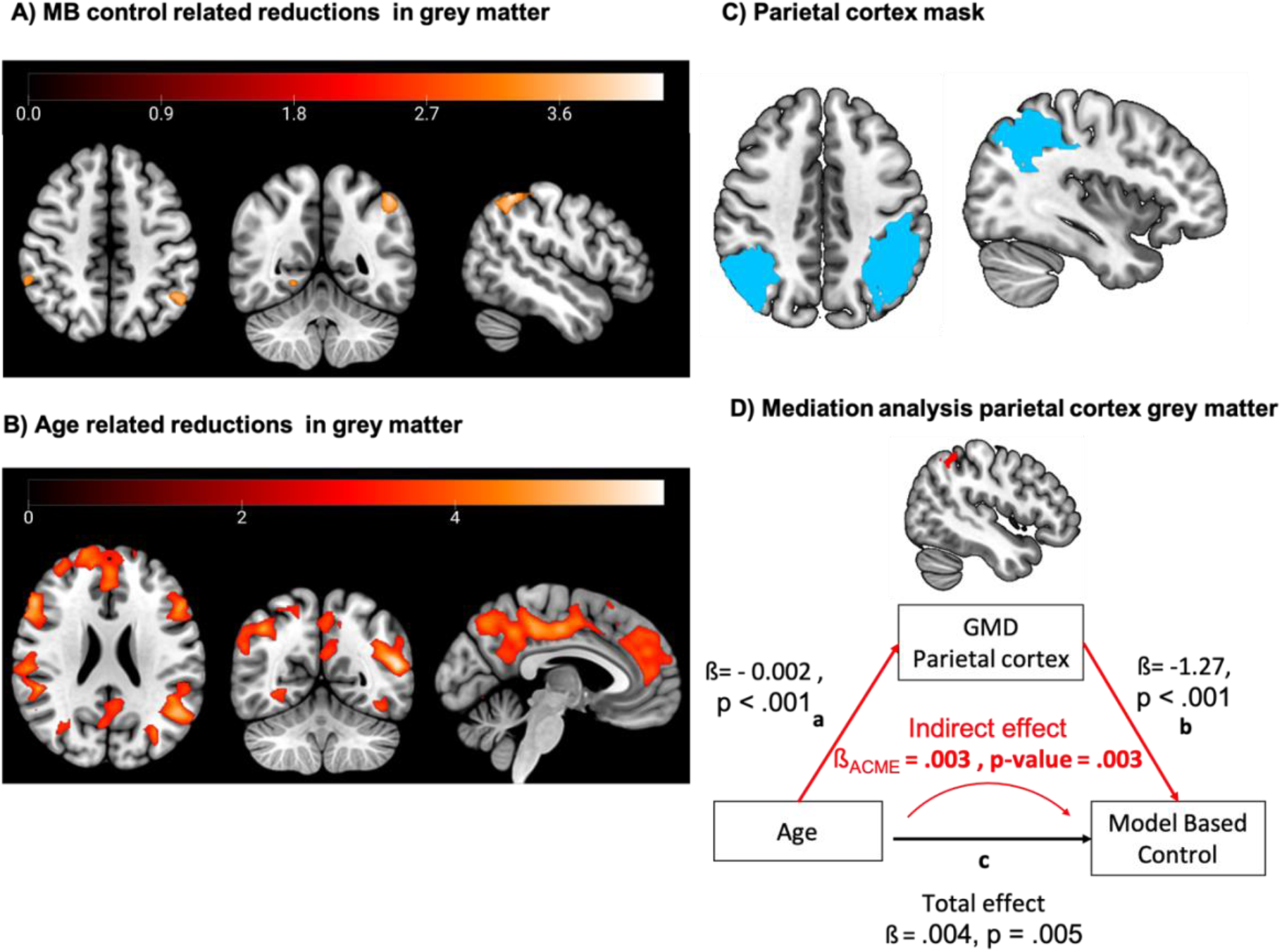
A) GM increases related to MB control in GLM 1 (map thresholded at voxel > 1099 depicting clusters significant at the whole brain level. B) Age related GM reductions found in the second GLM C) Depiction of parietal cortex mask based derived from the wfupicklas D) Mediation analysis showing the mediation effect of GMD extracted from an *a-priori* defined brain mask in the parietal cortex on the association between age and MB control.

### Learning from positive and negative feedback

Examining the ability to learn from positive and negative feedback using mixed-effect models, we found a significant main effect of learning from positive and negative feedback (ß = 0.321, SE = 0.1, χ^2^ = 10.73, p = .001), indicating that overall participants were better in choosing the best cue (learning from positive feedback) relative to avoiding the worst cue (Select best: 68.7% (23.2) vs. Avoid Worst 57.8% (23.2), Figure 3). Using correlational analysis, we found that learning from positive feedback improved with age (r_s_(88) = 0.35, p < .001), while the ability to learn from negative feedback did not (r_s_(88) = 0.07, p = .5) (c.f. Figure 3).

### MB control and feedback learning

MB control was positively correlated to positive feedback learning in the choice task (r_s_(88) = 0.28, p = .007). Negative feedback learning did not correlate significantly with MB control (r_s_(88) = 0.15, p = .2).

### Brain structural correlates of positive feedback learning

We then examined the neuroanatomical correlates of age-dependent increases in learning from positive feedback. In the GLM_PosFB_, we found positive feedback learning to be negatively related to GMD in two clusters on the whole brain level. Specifically, in the left and right frontal superior medial gyrus (vmPFC, MNI Peak coordinate: 9 69 22, k= 1037, T =5.02, p_FWE corr peak_= .02) and to GMD in a cluster in the right inferior and middle temporal gyrus (MNI Peak coordinate: 63 -60 -12, k= 446, T = 5.08, p_FWE corr peak_= .02). Using SVC, we also found a significant cluster in the parietal ROI (MNI Peak coordinate: -45 63 24, k= 71, T =4.12, p_FWE corr peak_= .02) and a significant cluster in the dlPFC ROI (MNI Peak coordinate: 20 63 26, k= 104, T =4.30, p_FWE corr peak_= .02).

In the GLM_PosFB_Full_, we saw widespread GMD changes that were negatively associated with age. For positive feedback learning, only one cluster in the supplemental motor area survived whole brain correction on the peak level (MNI Peak coordinate: -3 16 70, k= 390, T = 4.97, p_FWE corr peak_= .03) while the previously significant frontal cluster from the first GLM failed to reach significance, when controlling for age effects. We found significant clusters in the parietal ROI (MNI Peak coordinate: 60 -44 45, k= 197, T =4.21, p_FWE corr peak_= .02) and dlPFC (MNI Peak coordinate: -33 62 8, k= 546, T =4.72, p_FWE corr peak_= .005) using SVC, while clusters in the vmPFC did not survive multiple comparison correction. GMD changes were not significantly associated with the regressor for age x positive feedback learning.

We found the association for age and learning from positive outcomes to be partially mediated by GMD in all three regions of interest. GMD in the vmPFC accounted for up to 31.2 % of the total effect of the relationship of age and positive feedback learning (p = .046), while dlPFC GMD also mediated this association, explaining 42.4% (p = .01), as did GMD in the parietal cortex, explaining up to 57.9% (p = .01) of the effect (see Figure 4).

**Figure 4.**
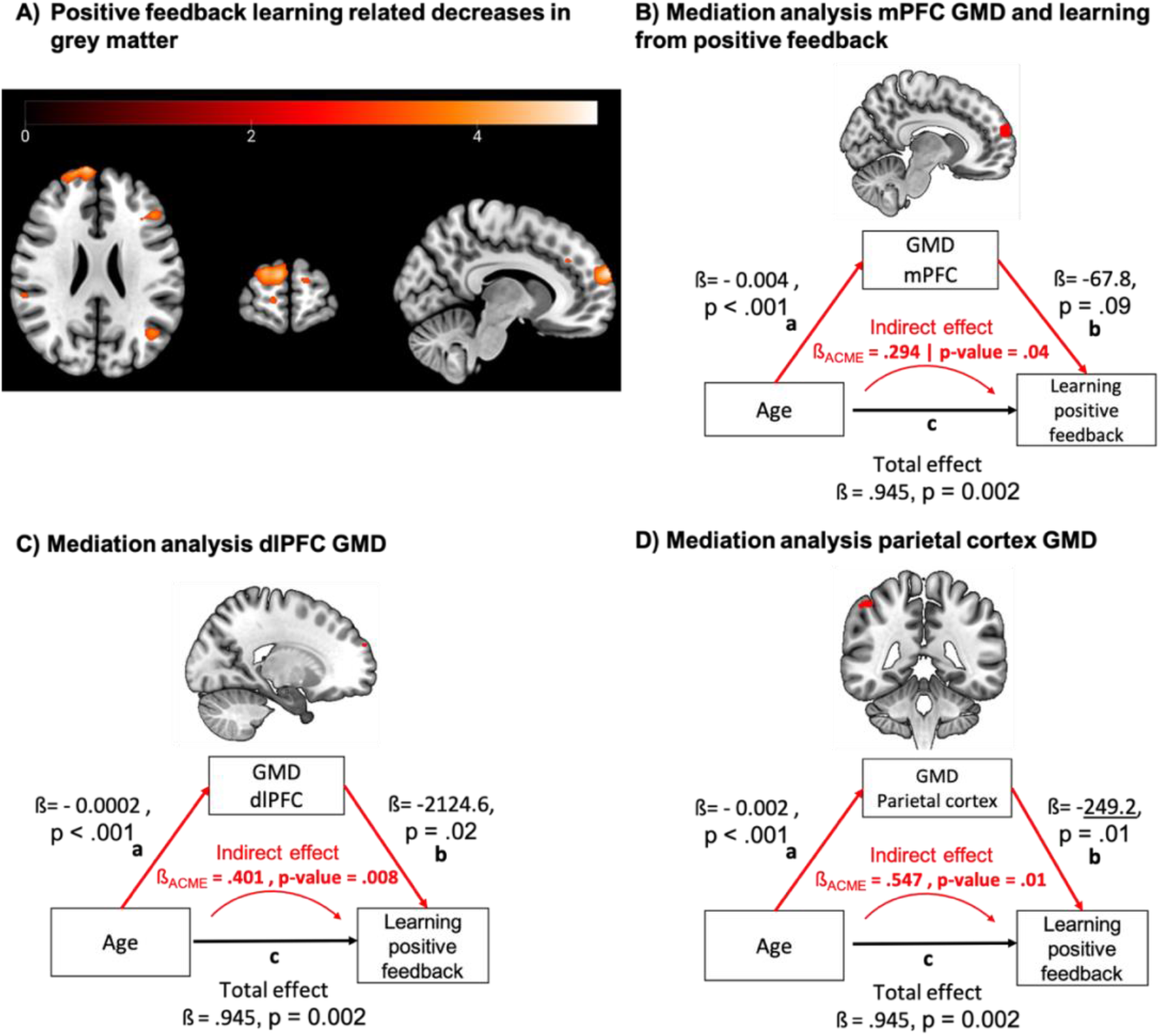
**A)** GMD reductions associated with increased learning from positive feedback as reported for the first GLM including MB control only as regressor. **B-D) Structural mediation analysis**. Depiction of significant mediation effect of GMD in a cluster in the mPFC, dlPFC and parietal cortex on the relationship between age and learning from positive feedback. P-values below .05 are considered significant.

## Discussion

In this study, we examined the brain structural correlates of age-dependent changes of MB control and feedback learning as a function of motivational context and outcome valence. We found age-related increases in MB control and positive feedback learning to be mediated by reduced GMD in the parietal cortex. Moreover, age effects for positive feedback learning were mediated by GMD in the vmPFC and dlPFC. Thus, we showed that previous findings of age-dependent increases of MB control are largely mediated by cognitive abilities. However, we found no evidence for motivational context relating to age-dependent changes in MB control. Finally, individuals relying more on MB control were also more adept at learning from positive feedback, suggesting improved value-based choices. Our findings thus provide significant new insight into the neuroanatomical correlates of age-related increases in MB control and positive feedback learning, while highlighting the need for considering motivational context and outcome valence within a developmental research framework.

To our knowledge, this is the first study examining the neuroanatomical correlates of developmental changes in MB control in different motivational contexts. Our finding of parietal GMD mediating a considerable portion (> 60%) of age effects on MB control corresponds to functional imaging work showing this region to encode the neural signature of MB control, a state prediction error (SPE), computing the difference between current state and observed state transition (Gläscher et al., 2010). It also agrees with the suggested role of posterior parietal regions as integration hub of spatial, temporal and reward information (Roitman et al., 2007).

We also find GMD in the parietal cortex, vmPFC and dlPFC to partially mediate age effects on positive feedback learning. The parietal cortex has been implicated in mapping the ordinal relationship among cues, a highly relevant feature when learning the relative rank order of task cues (Munoz et al., 2020). The vmPFC’s relevance for feedback learning is also supported by work tying mPFC and mOFC to adaptive decision making and subjective value encoding (Rushworth et al., 2011) for positive (Kringelbach, 2005) and negative outcomes (Tom et al., 2007). For the dlPFC, previous literature extensively highlighted its role in WM processes (Nee et al., 2013; Rottschy et al., 2012) further supported by lesion studies in humans (Barbey et al., 2013) and non-human primates (Butters et al., 1971; Butters & Pandya, 1969; Levy & Goldman-Rakic, 1999). Interestingly, one study showed a modulatory effect on reward and punishment sensitivity in a probabilistic RL task during non-invasive dlPFC stimulation possibly mediated through increased dopaminergic release in the ventral striatum (Ott et al., 2011). Thus, it is conceivable that the reported mediation effects of dlPFC GMD on the association between age and positive feedback learning is conveyed through WM processes facilitated in the dlPFC, which get updated in a feedback-dependent manner.

Previous developmental studies consistently reported age-dependent increases in MB control (Bolenz et al., 2017; Decker et al., 2016; Nussenbaum et al., 2020; Vaghi et al., 2020), but were not designed to disentangle motivational context effects. Interestingly, we see a lack of motivational context effects on MB control (and age-dependent effects therein), despite ample empirical evidence describing the influence of contextual effects on RL (Bavard et al., 2018; Louie & De Martino, 2014; Palminteri et al., 2015; Palminteri & Lebreton, 2021; Pischedda et al., 2020). This replicates a previous null finding of another study also examining motivational context effects in a developmental sample (Bolenz & Eppinger, 2021). This absence might be explained by the way motivational context can distinctively impact value updating depending on the reference point (Palminteri et al., 2015; Palminteri & Lebreton, 2021), which might have been facilitated by our block design. For illustration, in a punishment context, successful punishment avoidance (neutral feedback) can be perceived as rewarding, thus reinforcing the chosen option, while in a reward context, neutral feedback can be perceived as punitive, given the potential of gaining reward (Palminteri et al., 2015). It is thus conceivable that participants adjusted their reference point according to motivational context. Another explanation might arise from the employed task version, as more MB control does not translate into increased payouts or loss avoidance (Kool et al., 2016), potentially resulting in reduced sensitivity for motivational context. Still, this appears unlikely given previous work using a task variant addressing this short-coming, which also failed to report motivational context effects on MB control (Bolenz & Eppinger, 2021). Thus, while replicating previous findings of age-dependent increases of MB-control (Decker et al., 2016; Nussenbaum et al., 2020), we find no support for our hypothesis that adolescents are willing to exert relatively more effortful control by employing more MB control in a reward vs. punishment context.

Choice test performance indicated an age-dependent increase for positive feedback learning, while negative feedback learning seemed stable across development. Previous studies have already pointed to adults exhibiting better reward relative to punishment learning compared to adolescents (Palminteri et al., 2016; Van Der Schaaf et al., 2011) and increasing reward learning rates from childhood into adulthood suggest faster adaptation following rewards (Van Den Bos et al., 2012). In contrast, other studies reported age-dependent increases in negative learning rates, while positive learning rates remained stable (Rosenbaum et al., 2022, Pauli et al. 2022). Adolescents were inclined to focus more on worse-than-expected outcomes during learning and this tendency was associated with a subsequent memory recall bias (Rosenbaum et al., 2022). Interestingly, it has been proposed that the often-reported elevated reward responsivity in adolescents might not originate from enhanced reward learning but instead reflect a tendency of more pronounced action initiation during this phase (Pauli et al. 2022). Heterogenous findings might result from significant differences in task design but need to be further investigated to better understand the dynamics of age-dependent valence asymmetries in RL (Nussenbaum & Hartley, 2019). Overall, our results show that while recruitment of MB control is unaffected by motivational context, the way we learn and encode outcome-representations seems biased towards positive feedback learning. Interestingly, task performance across tasks was connected, with more MB control linked to better positive feedback learning. This suggest that independent of motivational context, more MB control might facilitate improved encoding of outcome-value representations for the respective cues in the first task phase allowing for better performance when selecting the ‘better’ cue during the choice task. This did not apply for negative feedback learning.

Interestingly, while the choice task was designed to distinguish MF and MB components, it has been posited that the asymmetric, valenced choice signature of feedback learning could be model-free (Collins & Frank, 2014; Doll et al., 2016; Frank, 2005). Thus, it might come as a surprise that the latter signature was linked positively to model-based control, at least when adhering to a strict dual system separation. Still, previous work indicated that such distinction might not be appropriate by showing that the MF system can have access to MB information (e.g. Daw et al., 2011; Deserno et al., 2021; Moran et al., 2019). Second, the association might arise from inherent experimental features given that our model-based index and the choice signature of feedback learning were derived from two highly interdependent tasks. It is also possible that the positive correlation between both variables reflects the ability to sharply represent values of distinct choice option, a feature that would be supportive for both MF and MB processes. Future studies are thus warranted to precisely characterize these task signatures.

Our data provides new insights in how brain structural changes impact the age-dependent maturation of decision processes. Unlike previous developmental studies, which primarily compared children and adolescents with adults below 30 years of age (Decker et al., 2016; Potter et al., 2017; Van Den Bos et al., 2012), our study fills an important gap in the literature by assessing individuals across a broad age range including middle adulthood. Our findings of increased MB control and feedback learning provide exciting new insights into decision and learning systems, both of which might have the capacity of further improving into middle adulthood. Interestingly, older individuals learned better from negative feedback relative to younger adults (Eppinger & Kray, 2011), suggesting that feedback learning in old age might not follow the trajectory seen in adolescence and adults (Ferreira et al., 2015; Singh-Manoux et al., 2012).

Regarding limitations, firstly, our study has a cross-sectional design and our findings are correlational in nature. More longitudinal studies (Vaghi et al., 2020) are needed to replicate our findings, while also tracking inter-individual developmental trajectories of the reported associations, which can otherwise go unnoticed in cross-sectional design (Koolschijn et al., 2011). This seems especially important considering substantial variations in structural brain development across subcortical and cortical regions (Mills et al., 2021; Wierenga et al., 2014). Secondly, the use of different imaging sequences is not desirable but was controlled for.

In sum, our study shows that age-related changes of MB-control and positive feedback learning are mediated by GMD in the parietal cortex, dlPFC and vmPFC. Meanwhile, more MB control also translates into improved selection of cues linked to better outcomes. Our results underline the importance of taking into account valence effects by teasing apart the role of 1) motivational context and 2) learning from positive and negative outcome when studying decision and learning processes in the framework of developmental research. The potential implications of our findings for neurodevelopmental disorders such as ADHD and compulsivity, including how developmental trajectories of MB control and feedback learning might diverge in non-healthy development, should be the focus of future longitudinal studies.

## Code and Data availability

The anonymized behavioral raw data and 2^nd^ level models from the analysis of structural brain data for this study will be shared via an online repository.

## Author contributions

L.D., A.M.F.R. and AH discussed initial ideas for the study. L.D., A.H., N.H. and M.W. were responsible for designing and implementing the study. M.W. and N.H. collected the data. V.S. and L.D. analyzed the data. V.S. and L.D. wrote the manuscript. All authors were involved in interpreting the results and were responsible for reviewing and editing the manuscript.

## Acknowledgements

This work was directly funded by a Grant to L.D. and A.H. from the IFB Adiposity Diseases, Federal Ministry of Education and Research (BMBF), Germany, GN: 01EO150. LD also received funding from the German Research Foundation (DFG) as part of the Collaborative Research Centre 265 “Losing and Regaining Control over drug intake” (Project A02), which partially supported this work. A.M.F.R. acknowledges support from grants by the German Research Foundation (Deutsche Forschungsgemeinschaft, DFG RE 4449/1-1, SFB 940-3/B7, RTG-2660/B2) and by a 2020 BBRF Young Investigator Grant. The funders have no role in study design, data collection and analysis, decision to publish or preparation of the manuscript.

